# Breaking the ladder: Evolution of the ventral nerve cord in Annelida

**DOI:** 10.1101/378661

**Authors:** Conrad Helm, Patrick Beckers, Thomas Bartolomaeus, Stephan H. Drukewitz, Ioannis Kourtesis, Anne Weigert, Günter Purschke, Katrine Worsaae, Torsten H. Struck, Christoph Bleidorn

**Affiliations:** Animal Evolution and Biodiversity, Georg-August-University Göttingen, 37073 Göttingen, Germany; Institute of Evolutionary Biology and Ecology, University of Bonn, 53121 Bonn, Germany; Institute of Biology, University of Leipzig, 04103 Leipzig, Germany; Department of Evolutionary Genetics, Max Planck Institute for Evolutionary Anthropology, 04103 Leipzig, Germany; Department of Developmental Biology and Zoology, University of Osnabrück, 49069 Osnabrück, Germany; Department of Biology, University of Copenhagen, 2100 Copenhagen, Denmark; Frontiers in Evolutionary Zoology, Natural History Museum, University of Oslo, P.O. Box 1172, Blindern, NO-0318 Oslo, Norway; German Centre for Integrative Biodiversity Research (iDiv) Halle-Jena-Leipzig, Deutscher Platz 5e, 04103 Leipzig

## Abstract

A median, segmented, annelid nerve cord has repeatedly been compared to the arthropod and vertebrate nerve cords and became the most used textbook representation of the annelid nervous system. Recent phylogenomic analyses, however, challenge the hypothesis that a subepidermal rope-ladder-like ventral nerve cord (VNC) composed of a paired serial chain of ganglia and somata-free connectives represents neither a plesiomorphic nor a typical condition in annelids.

Using a comparative approach by combining phylogenomic analyses with morphological methods (immunohistochemistry and CLSM, histology and TEM), we compiled a comprehensive dataset to reconstruct the evolution of the annelid VNC. Our phylogenomic analyses generally support previous topologies. However, the so far hard-to-place Apistobranchidae and Psammodrilidae are now incorporated among the basally branching annelids with high support. Based on this topology we reconstruct an intraepidermal VNC as ancestral state in Annelida. Thus, a subepidermal ladder-like nerve cord clearly represents a derived condition.

Based on the presented data, a ladder-like appearance of the ventral nerve cord evolved repeatedly, and independently of the transition from an intraepidermal to a subepidermal cord during annelid evolution. Our investigations thereby question a common origin of the bilaterian median ganglionated VNC and propose an alternative set of neuroanatomical characteristics of the last common ancestor of Annelida or perhaps even Spiralia.

## Background

A rope-ladder-like organization of the ventral nerve cord (VNC) has often been regarded to represent the ancestral condition of annelids [1–7]. According to this traditional view, the VNC in Annelida consists of a chain of paired ganglia containing the neuronal somata, linked longitudinally by parallel somata-free connectives and transversely by segmental commissures [1–4, 7]. The organization, development and cell types of the annelid mid-ventral cord are best investigated in the annelid model organisms *Capitella teleta* Blake, Grassle & Eckelbarger, 2009, *Helobdella robusta* Shankland, Bissen & Weisblat, 1992 and *Platynereis dumerilii* (Audouin & Milne Edwards, 1834) [8–15]. Gene expression studies in these annelids inspired wide reaching comparisons of the annelid VNC to that of arthropods and vertebrates (e.g., [16–19]). Nonetheless, the annelid VNC shows a great diversity in number and position of neurite bundles and localization either within or beneath the epidermis [1, 4, 5, 7, 20, 21]. Accordingly, the hypothesis that the ladder-like VNC is ancestral was often questioned [21–24] and challenged repeatedly, e.g., by a hypothesis regarding a pentaneuralian arrangement of the neurite bundles in the annelid VNC as ancestral [4, 5; rejected in 25] and by the finding of an unpaired mid-ventral nerve cord in numerous taxa [15, 21, 24, 26–28]. Recent, well-supported phylogenomic analyses [29–33] revealed that previous profound investigations into annelid neuroanatomy unfortunately focussed on representatives of derived annelid subgroups, now united as Pleistoannelida [4, 5, 21]. However, several taxa placed outside this main clade have been neglected so far. For instance, Magelonidae and Oweniidae together (as Palaeoannelida) represent the sister taxon of all other annelids. Subsequently, b Chaetopteridae and a clade comprising Sipuncula and Amphinomida are branching of. Comparative neuroanatomical investigations focussing on these non-pleistoannelid taxa are still limited [34–38]. Moreover, several groups that were so far difficult to place in the annelid tree but were sometimes also considered as possibly early-branching, namely Apistobranchidae and Psammodrilidae, were neither included into recent phylogenomic studies [39, 40] nor examined in detail concerning their neuroanatomy [41, 42].

In order to understand the VNC evolution in Annelida we investigated this character complex within these so far neglected non-pleistoannelidan taxa. Therefore, we examined the trunk nervous system of 19 taxa with focus on the position of the VNC within or outside the epidermis, the arrangement and immunoreactivity of neurite bundles within the VNC, the appearance of additional neurites such as giant fibers as well as shape, arrangement and location of neuronal somata along the VNC. Further special attention was given to presence or absence of somata-free connectives and commissures along the entire VNC. Moreover, we updated previous phylogenomic datasets with the hitherto neglected groups Apistobranchidae and Psammodrilidae. Combining these transcriptomic analyses with immunohistochemistry and confocal laser scanning microscopy (CLSM), histological Azan staining and transmission electron microscopy (TEM), we compiled an updated phylogenomic tree and a comprehensive neuroanatomical dataset. Our results provide the background for a better understanding of the nervous system evolution within Annelida and Spiralia in general.

## Methods

### Collection and fixation of specimens

We collected fresh specimens of several species for our study, representing 14 annelid families: 10 species solely for RNA extraction and subsequent transcriptomic analyses and 24 species for morphological investigations. For further details, please refer to additional table S1. Divergent collection and fixation details are specified were required and references are given below.

### Transcriptome library construction and Illumina sequencing

RNA extraction and library construction were conducted as described in detail in Weigert et al. [30]. Newly constructed libraries, as well as several libraries only shallowly covered in previous analyses [30], were sequenced on an Illumina HiSeq 2500 100 bp paired-end. Base calling was performed with freeIBIS [43], adaptor and primer sequences were removed, low complexity reads and false paired indices were discarded. Raw data of all libraries were trimmed by applying a filter of Phred 15. Data for additional taxa were obtained from NCBI (National Center for Biotechnology Information (NCBI) run by the National Institutes of Health) (see supplementary material table S2). Libraries were assembled *de novo* using either the CLC Genomics Workbench 5.1 (CLC bio, Århus, Denmark) or Trinity [44].

### Phylogenomic analyses

A list of all taxa used and the source of data is given in additional file table S2. Orthology prediction was performed using HaMStR [45]. The applied core-orthologs set comprises 1,253 orthologous genes downloaded from the Inparanoid database [46]. *Capitella teleta, Helobdella robusta, Lottia gigantea, Schistosoma mansoni, Daphnia pulex, Apis mellifera*, and *Caenorhabditis elegans* served as primer-taxa. Redundant sequences were eliminated using a custom Perl script [30].

Alignments for each orthologous gene were generated separately using MAFFT [47] and alignment masking was performed with REAP [48]. All masked single gene alignments were concatenated into a supermatrix using a custom Perl script. To reduce potential problems of missing data we compiled two data matrices using the program MARE [49] with weighing parameters of α=1.5 and α=2, resulting in two differently densely covered supermatrices. We used partition finding and model testing as implemented in IQ-TREE [50] for both supermatrices, subsequently analysed under the Maximum Likelihood optimality criterion as implemented in the same program. Bootstrap support was estimated from 1000 pseudoreplicates.

### Azan staining, histological sections and 3D-reconstruction

Adult specimens were fixed (see additional file table S1 for species details), stained and analyzed as described in Beckers et al. [52]. Thus, the specimens were fixed overnight in Bouin’s fixative modified after Dubosque-Basil, dehydrated in an ethanol series and incubated in methylbenzoat and butanol. Afterwards the samples were preincubated in Histoplast (Thermo Scientific, Dreieich, Germany) and embedded in Paraplast (McCormick Scientific, Richmond, USA). 5 μm thick sections were made using a Reichert-Jung Autocut 2050 microtome (Leica, Wetzlar, Germany) and transferred to albumen-glycerin coated glass slides. Sections were stained with Carmaulaun, differentiated with sodium phosphotungstate (5%), washed in distilled water, stained in aniline blue orange G and subsequently embedded with Malinol (Waldeck, Münster, Germany). In Azan staining, the neuropil of the nervous system stains gray, the nuclei of cell somata stain red, the extracellular matrix stains blue and the musculature stains orange [52]. Each section was digitalized at 40x magnification using a slide scanner (Olympus dotslide (2.2 Olympus, Hamburg) and aligned using IMOD [53] and imodalign (http://www.qterra.de/biowelt/3drekon/guides/imod_first_aid.pdf). 3D reconstructions were performed with Fiji (1.45b) [54], trakem [55] and Amira (4.0).

### Ultra-thin sections and transmission electron microscopy (TEM)

For electron microscopy animals were either fixed in 1.25% glutaraldehyde buffered in 0.05 M phosphate buffer containing 0.3 M NaCl for 1 hour, rinsed several times in the same buffer and postfixed in 1% OsO4 buffered in the same manner (for *Owenia fusiformis, Magelona mirabilis, Spiochaetopterus costarum, Chaetopterus variopedatus* and *Psammodrilus balanoglossoides*), in 2.5 % glutaraldehyde/ 0.1 M sodium cacodylate/ 0.24 M NaCl and subsequently post-fixed in 1 % OsO4/ 0.1 M sodium cacodylate/ 0.24 M NaCl (for *Apistobranchus tullbergi*) or in a phosphate-buffered mixture of sucrose, picric acid, glutaraldehyde and paraformaldehyde (SPAFG), according to Ermak and Eakin [56], for 2.5 hours at 4°C and rinsed in 0.075 M phosphate buffer adjusted to seawater with sucrose (7 changes, 2 hours) (for *Eurythoe complanata* and *Paramphimone* sp.). In the latter case specimens were postfixed in 1% OsO4 in the same phosphate buffer for 1 h at 4°C, otherwise the specimens were stained for 30 min in 2 % OsO4/ 1.5 % potassium ferricyanide/ 0.1 M sodium cacodylate followed by incubation in 2 % aqueous uranyl acetate for 30 min. Dehydration of the samples was performed gradually in a graded ethanol or an ascending acetone series and then with propylene oxide. All steps were conducted at room temperature. Following embedding (using the TAAB Araldite 502/812 kit or Epon-Araldite 812 kit), ultrathin sections (70 nm) were cut with a Leica Ultracut E, UC6 or UC7 and counterstained with 2 % uranyl acetate and lead citrate. Images were acquired on JEOL 1011, Zeiss EM 902A, Zeiss EM 10CR, Zeiss Lyra or Zeiss Libra 120 transmission electron microscopes equipped with an Olympus MORADA or a 4K TRS (Moorenweis, Germany) camera. Figures were adjusted to 8-bit grey scaling with the Analysis software package. All final panels were prepared using Adobe (San Jose, CA, USA) Photoshop CC and Illustrator CC.

### Immunohistochemistry, CLSM and image processing

Although the specificities of the employed antibodies have all been established in numerous invertebrates, we cannot fully exclude that a given antiserum may bind to a related antigen in the investigated specimens. We hence refer to observed labelled profiles as exhibiting antigen-like immunoreactivity (–LIR). For subsequent staining, at least five specimens of each taxon where used (see electronic supplementary material Table S1 for details). Antibody staining was preceded by tissue permeabilisation for at least 1 h in 0.1 M PBS containing 0.1% NaN3 and 0.1% TritonX-100 (PTA), suited by blocking in block-PTA (6% normal goat serum (Sigma-Aldrich, St. Louis, MO, USA) in PTA) for 2-4 hours or overnight. The primary antibodies, polyclonal rabbit antiserotonin (INCSTAR, Stillwater, USA, dilution 1:500), polyclonal rabbit anti-FMRFamide (Acris Antibodies GmbH, Herford, Germany, dilution 1:500) and monoclonal mouse anti-acetylated α-tubulin (clone 6-11B-1, Sigma-Aldrich, St. Louis, USA, dilution 1:500) were applied for 48-72 hours in block-PTA. Afterwards, specimens were rinsed in block-PTA for 3 × 2 hours and incubated subsequently with secondary fluorochrome conjugated antibodies (goat anti-rabbit Alexa Fluor 488) in block-PTA for 24-48 hours. At last, the samples were washed three times in 0.1 M PBS (without NaN3). Subsequently the samples were mounted between two cover slips using 90% glycerol/ 10% 10x PBS containing DABCO or Vectashield Mounting Medium (Vector Laboratories, Burlingame, USA) (for *Apistobranchus*). Negative controls were obtained by omitting the primary antibody in order to check for antibody specificity and yielded no fluorescence signal. Specimens were analyzed with the confocal laser-scanning microscope Leica TCS STED (Leica Microsystems, Wetzlar, Germany). Confocal image stacks were processed with Leica AS AF v2.3.5 (Leica Microsystems). The final panels were designed using Adobe (San Jose, CA, USA) Photoshop CC and Illustrator CC.

### Character definition

Ten morphological characters were defined as primary homologies in order to reconstruct the ancestral state of the annelid ventral nerve cord. All characters are defined as binary and their codings and relevant references shown in the data matrix of figure 6. Codings are based on the newly generated data. Inapplicable character states are treated as missing data in our analyses.

#### Character 1: Intraepidermal position of the ventral nerve cord

The ventral nerve cord of Annelida can be located within the epidermis (= intraepidermal, state: 1) or outside the epidermis (= subepidermal, state: 0). Intraepidermal cords are embedded within epidermal tissue and surrounded by epidermal cells. The basal lamina delimits the epidermis (including the ventral nerve cord) from the remaining non-epidermal tissue.

#### Character 2: Clusters of somata along the ventral nerve cord

Somata of the ventral nerve cord can be clustered (state: 1) or non-clustered (state: 0). Non-clustered somata don’t show any sign of aggregation, are evenly distributed along the nerve cord, and a neuropil is present along the entire nerve cord.

#### Character 3: Somata between clusters along the ventral nerve cord

If the somata along the ventral nerve cord are clustered according to character 2, single somata can appear between these clusters (state: 1). The latter somata are not part of the cluster, but belong to the ventral nerve cord. If somata between the clusters are absent (state: 0), these inter-clustal parts of the ventral nerve cord are called somata-free connectives. If the somata in character 2 were scored as non-clustered, they would here be scored as inapplicable.

#### Character 4: Segmentally arranged commissures along the ventral nerve cord

The commissures along the ventral nerve cord, interconnecting both parallel cords, can be arranged in a quite random series (state: 0) or in a segmental pattern (state: 1), congruent to the serial repetition of other structures. In both cases faint and prominent commissures are counted with the same weight.

#### Character 5: Different morphotypes of somata along the ventral nerve cord

The somata forming the ventral nerve cord can be monomorphic (state: 0) or polymorphic (state: 1).

#### Character 6: Giant fibers in the ventral nerve cord

Within the ventral nerve cord giant fibers can be absent (state: 0) or present (state: 1). These fibers are characterized by their larger diameter in comparison to other neurites forming the ventral nerve cord and run along the entire length of the ventral nerve cord.

#### Character 7: Ventral position of giant fibers within the ventral nerve cord

Giant fibers within the ventral nerve cord can be exclusively located in a ventral position (state: 1), or they can vary in their localization (state: 0). If giant fibers are absent, the character is coded as inapplicable.

#### Character 8: Number of giant fibers in the ventral nerve cord

Giant fibers can be present as a pair (state: 0) or be numerous (state: 1). If giant fibers are absent, the character is coded as inapplicable.

#### Character 9: Fusion of giant fibers in the ventral nerve cord

Giant fibers in the ventral nerve cord can be separate (state: 0) or partly fused (state: 1) throughout the trunk. If giant fibers are absent, the character is coded as inapplicable.

#### Character 10: Additional longitudinal neural bundles originating in the brain

In close proximity to the ventral nerve cord additional longitudinal neurites can be present (state: 1) or absent (state: 0). Notably, these additional longitudinal neurites arise from the brain and do not branch off from the ventral nerve cord.

### Ancestral state reconstruction

Ancestral states for separate characters of the ventral nerve cord were reconstructed in Mesquite v. 3.10 [51]. In a first analysis a parsimony approach with characters treated as unordered was used. A second analysis was carried out using a maximum likelihood reconstruction showing proportional likelihoods under the Mk1 model with branch lengths scored as equal. Both reconstructions were based on a simplified topology of the MARE1.5-tree (figure 1), the underlying topology was predefined by hand and potential annelid sister groups were not included due to the unresolved annelid sister group. Results of both analyses were highly similar and only the maximum parsimony reconstruction is shown in the main text. A summary of this reconstruction is shown in figure 7, and the actual ancestral state reconstructions (for both methods and each character) are provided as additional files S4 (MP reconstruction) and S5 (ML reconstruction). Reconstruction of the characters on the MARE 2.0-tree (figure S3) were not shown, but the only discrepancy found (character 6) was discussed in the text.

**Figure 1.**
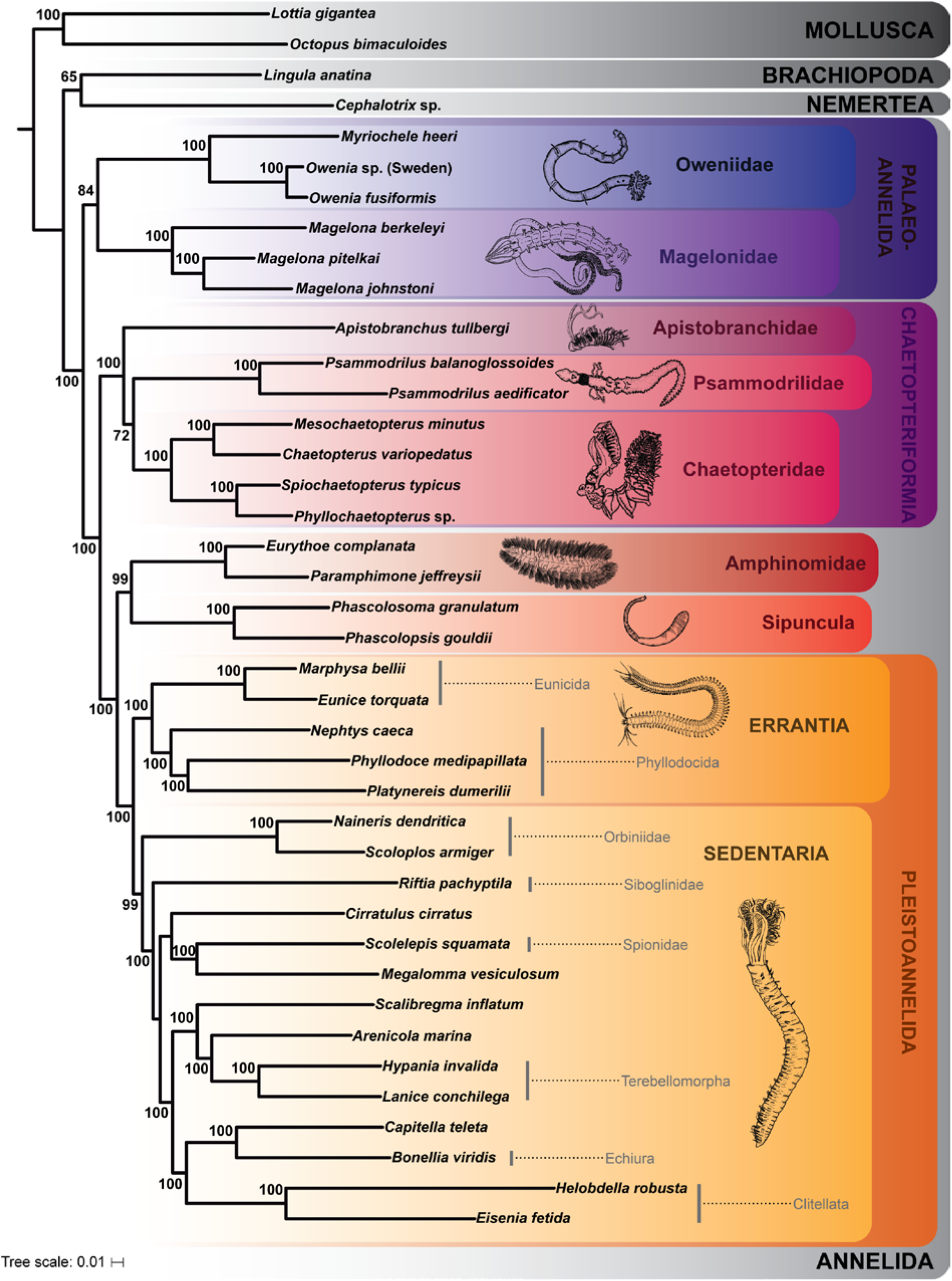
Best maximum likelihood (ML) tree of the RAxML analysis using the MARE1.5 data set of 40 taxa, including 490 gene partitions comprising 159,297 amino acid positions. Only bootstrap values above 50 are shown. The drawings are redrawn from various sources.

To test for the influence of potential annelid sister groups on the ancestral state reconstruction, MP analyses were also performed with either Mollusca, Nemertea, Phoronida or Brachiopoda as potential annelid sister groups. The detailed results are given in the additional files S6 (for Mollusca), S7 (for Nemertea), S8 (for Phoronida) and S9 (for Brachiopoda).

## Results and Discussion

### Molecular analyses

Two supermatrices were used for phylogenetic analysis, which differed in the number of included gene partitions. Using the weighting parameter α=2 resulted in a matrix containing 404 of the original 1,253 gene partitions, comprising 128,186 amino acid positions for the 40 analyzed taxa (MARE2 dataset). Applying the less strict weighting parameter α=1.5 kept 490 partitions for the final supermatrix (MARE1.5 dataset), comprising 159,297 amino acid positions. With 90.1% (MARE2) and 90.6% (MARE1.5), the completeness of the matrix is similar for both data sets and clearly improved compared to the unreduced supermatrix (24.2%). Maximum Likelihood analyses using a partition scheme and models as optimized by IQ-TREE [50] resulted in two slightly different topologies (figures 1, S3).

Consistent with previous analyses [29, 30, 33], both data sets yielded a monophyletic Annelida, which can be broadly classified into Pleistoannelida (comprising Sedentaria and Errantia) as well as a number of basally branching lineages (see figure 1). As in previous analyses (see [33]), Sipuncula + Amphinomida are recovered as sister taxon to Pleistoannelida. Interestingly, we find Apistobranchidae and Psammodrilidae together with Chaetopteridae in a well-supported (100% bootstrap) clade among the basally branching lineages. Accordingly, the clade is herein named ‘Chaetopteriformia’ due to the well-supported placement of Apistobranchidae and Psammodrilidae together with Chaetopteridae. ‘Formia’ is derived from ‘forma’, which means ‘shape’, and follows previous naming for groups of annelid families such as Aphroditiformia, Cirratuliformia, Terebelliformia [see e.g., 30]. The naming is so far node-based and further morphological investigations (including the herein presented data) will help to provide a proper description and test for morphological apomorphies of this group in the future. Apistobranchidae and Psammodrilidae had not been included in previous phylogenomic analyses of annelid relationships and previous morphological investigations suggested them being positioned within Sedentaria mainly based on the structure of the palps, coelomic cavities or chaetal characters [57–60]. Due to the lack of detailed morphological investigations, morphological characters supporting the clade Chaetopteriformia are missing so far. Nevertheless, characters such as the arrangement of muscle bundles in the body wall, the presence and shape of internal chaetae or the partitioning of the trunk (heteronomous segmentation) should be analyzed in detail in future studies as they are candidates for putative morphological synapomorphies supporting the monophyly of Chaetopteriformia (see also [57] for review). In our analysis Psammodrilidae represent the sister taxon of Chaetopteridae, but the bootstrap support for this hypothesis is rather weak (bootstrap 72% & 78%, respectively). Nevertheless, the clade Chaetopteriformia and its position as sister group of (Sipuncula + Amphinomida) + Pleistoannelida (Errantia + Sedentaria) is maximally supported (figures 1, S3).

The two analyses only differ with respect to the sister taxon of all other annelids. In the MARE1.5 analyses (which contains more characters) Magelonidae and Oweniidae form the clade Palaeoannelida [33], which constitutes the sister group of all remaining annelids (see figure 1). This result is consistent with previous phylogenomic studies [30, 32, 61]. In contrast, in the MARE2 analysis, Oweniidae and Magelonidae branch of successively in the ML tree (additional file figure S3). However, in the bootstrap analysis the support for this latter topology is low, whereas the alternative hypothesis Palaeoannelida received higher support (bootstrap 84%). Therefore, we focused on the topology recovered by the MARE1.5 analysis for reconstructing the evolution of the VNC (figure 7), but discuss the few discrepancies found from the reconstructions performed on the MARE2-tree.

### Architecture of the VNC in early branching annelid taxa

In the following we refer to Richter et al. [62] regarding the terminology of neuroanatomical characters. Differing definitions are stated appropriately. Neuroanatomical characteristics of the VNC were observed in whole mounts of at least 5-10 adult specimens per species via immunohistochemistry and verified using histological series of sections and TEM in representative trunk regions. In particular, the position of the VNC in relation to the epidermal extracellular matrix (ECM), the arrangement and location of neurite bundles within the VNC and their respective immunoreactivity as well as the morphology and arrangement of neuronal somata in the VNC and the occurrence of somata-free areas were scored. Although different species per family were investigated and the coverage of morphological diversity within a family was given priority, it has to be kept in mind that the taxon sampling represents only a subset of the diversity within the respective families.

Our study revealed an intraepidermal VNC for all investigated species of Oweniidae (figures 2a, b, e), Magelonidae (figures 2g, h), Apistobranchidae (figures 3a, b), Psammodrilidae (figures 3f, h) and Chaetopteridae (figures 4a-c). These findings are concordant with previous investigations [34, 37, 42, 63, 64]. In species of Amphinomidae, the anterior part of the ventral cord is intraepidermal (figure 4g), whereas the posterior part is located subepidermally within the musculature (figure 4h, i), but still surrounded by a continuous ECM connected with the epidermal ECM (figure 4i).

**Figure 2.**
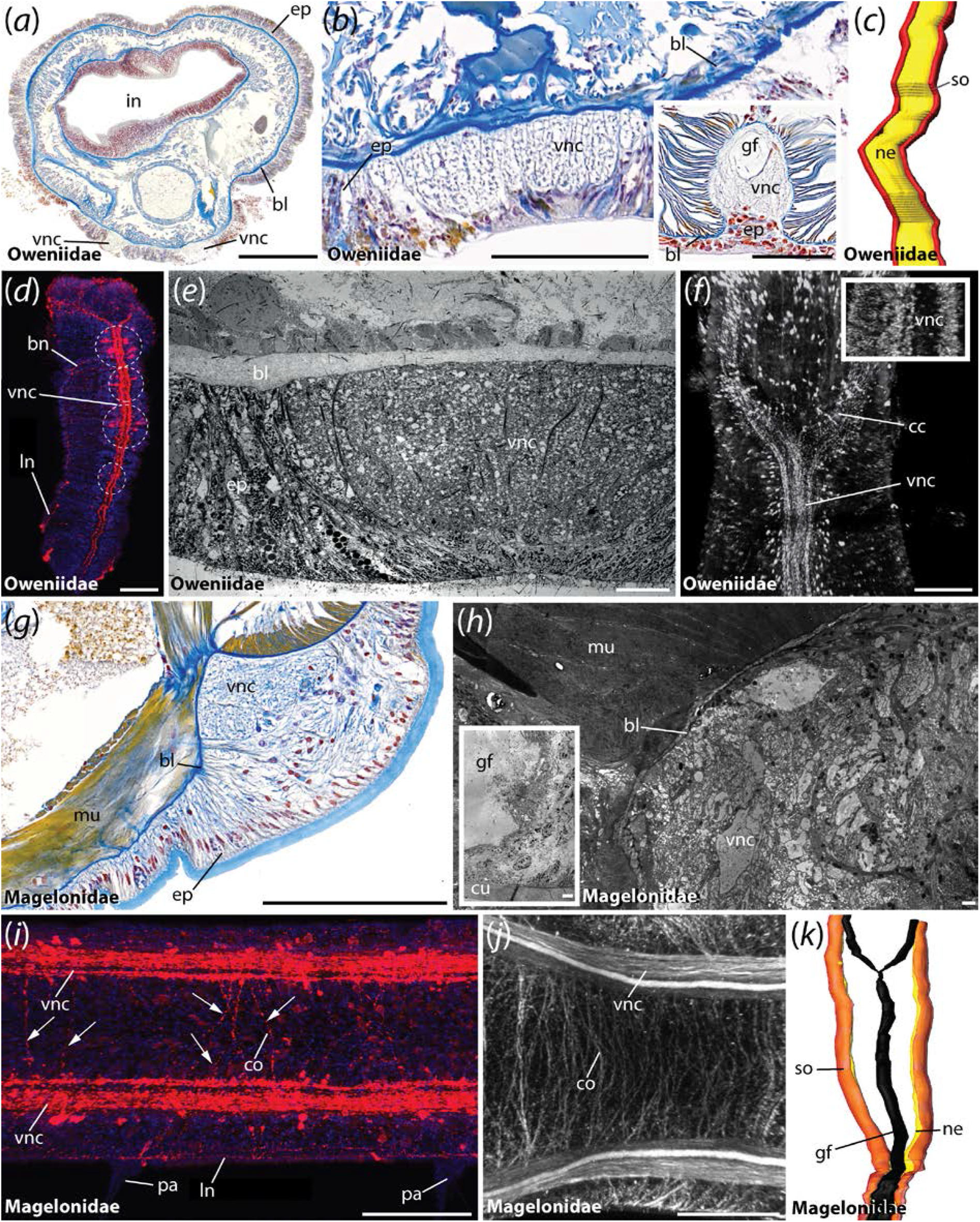
The VNC of Oweniidae and Magelonidae. Cross-sections (a, b, e, g, h) and ventral view (c, d, f, i-k) of the ventral cord in *Owenia fusiformis* (a-e), *Myriochele heeri* (f), *Myriowenia* sp. (b, iset), *Magelona mirabilis* (g, h, k) and *Magelona filiformis* (I, j). Anterior is up in (c, d, f, k) and left in (I, j). Azan staining (a, b, g), TEM (e, h), immuno-staining against 5-HT (d, f, i), tubulin (j) and DAPI (f, inset), and 3D-reconstruction of serial histological sections (c, k). (a, b) The VNC (vnc) in Oweniidae is situated within the epidermis. Giant fibers (gf) are only reported for Myriowenia (inset). (c) Oweniids show a medullary arrangement of somata (so) and neuroli (ne) along the ventral cord. No somata-free areas are detectable. (d) Juvenile oweniids exhibit serial immunoreactive clusters. (e) Ultrastructural data reveals a position of the ventral nerve cord (vnc) within the epidermis (ep). A basal lamina (bl) is present underlying the epidermis (ep). (f) anti-5-HT staining reveals a ventral nerve cord (vnc) with appearing immunoreactive somata in the anterior region. A DAPI staining in the same region reveals the lack of somata-free areas along the (vnc). (g) In magelonids the ventral cord (vnc) is comprised of two parallel cords in anterior segments, whereas the posterior ones only possess one fused (vnc). (h) Ultrastructural data verify the intraepidermal position of the (vnc) and the presence of giant fibers (gf). A basal lamina (bl) demarcates the subepidermal musculature (mu) from the intraepidermal nervous system (vnc). (i) Throughout the trunk the arrangement of somata can be regarded as being irregular. Numerous commissures are present (arrowheads). (j) Anti-tubulin staining reveals two parallel neurite bundles forming the ventral cord (vnc) in the trunk of adults. Furthermore, numerous commissures (co) are assembled along the medullary ventral nerve cord (vnc). (k) 3D-reconstructions verifies the fusion of the two parallel neurite bundles in the trunk of magelonids. bl, basal lamina; bn, branching nerve; cc, circumesophageal connective; co, commissure; cu, cuticle; ep, epidermis; gf, giant fibers; in, intestine; ln, longitudinal nerve; mu, muscle; ne, neuropil; pa, parapodia; so, somata; vnc, ventral nerve cord. Scale bars = 100 μm (a, b, d, I, j, f), 50 μm (b (inset), g), 10 μm (e) and 2.5 μm (h).

**Figure 3.**
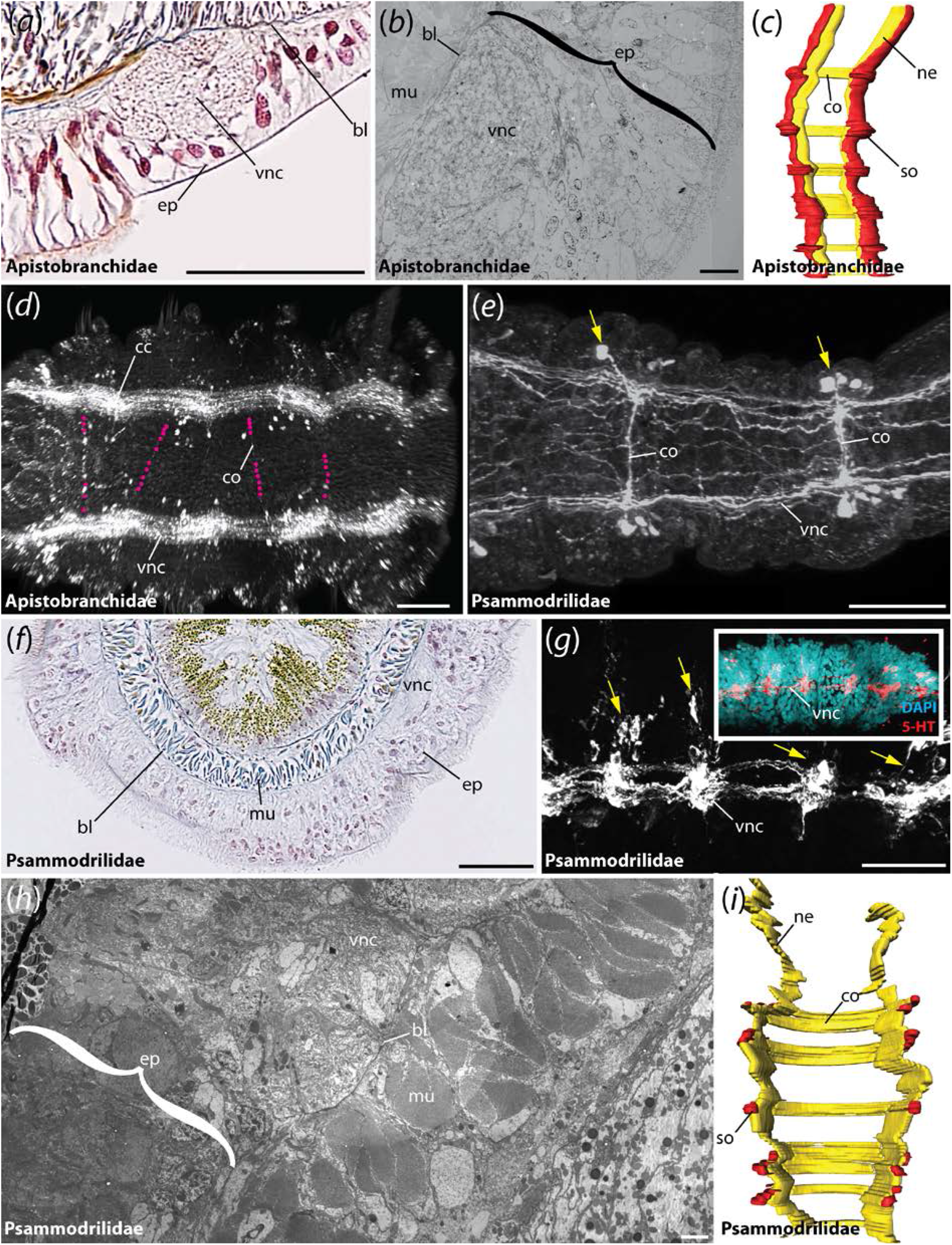
The VNC of Apistobranchidae and Psammodrilidae. Cross-sections (a, b, f, h), ventral (c-e, i) and lateral view (g) of the ventral cord in *Apistobranchus tullbergi* (a-d), *Psammodrilus balanoglossoides* (f, h, i) and *Psammodrilus curinigallettii* (e, g). Anterior is up in (c, i) and left in (d, e, g). Azan staining (a, f), TEM (b, h), immuno-staining against FMRFamide (d), 5-HT (e, g), 5-HT and DAPI (g, inset), and 3D-reconstruction of serial histological sections (c, i). (a) The VNC (vnc) in Apistobranchidae is situated within the epidermis (ep). The basal lamina (bl) demarcates the epidermal from the subepidermal tissue. Giant fibers are absent. (b) Ultrastructural data reveals a position of the ventral nerve cord (vnc) within the epidermis (ep). A basal lamina (bl) is present underlying the epidermis (ep). (c) Apistobranchidae show a medullary arrangement of somata (so) and neuropil (ne) along the ventral cord. Somata-free areas are absent. Nevertheless, the commissures (co) show a serial arrangement. (d) FMRFamide-immunoreactivity reveals the presence of two neurite bundles within the ventral nerve cord (vnc). Faint commissures (co) are visible as well. Note that for a better visibility the location of the commissures (co) is marked with purple dots. (e) Anti-5-HT staining of the ventral cord in Psammodrilidae reveals the presence of clustered somata (yellow arrowheads) and serially-arranged commissures (co). (g) The VNC (vnc) is situated within the epidermis (ep) as well. The underlying basal lamina (bl) is clearly visible. (g) Lateral view of the (vnc) verifies the presence of immunoreactive somata clusters (yellow arrowheads). Additional DAPI staining supports the anti-5-HT staining. (h) Ultrastructural investigations support the localization of the (vnc) within the epidermis (ep) and show the underlying basal lamina (bl) that demarcates epidermal and subepidermal layers. The musculature (mu) is situated in subepidermal position. (i) A 3-D reconstruction of serial histological sections in Psammodrilidae reveals the presence of clustered somata (so) along the ventral nerve cord. Serially arranged commissures (co) are present as well. bl, basal lamina; cc, circumesophageal connective; co, commissure; ep, epidermis; mu, muscle; ne, neuropil; pa, parapodia; so, somata; vnc, ventral nerve cord. Scale bars = 50 μm (a, e-g), 10 μm (b), 100 μm (d) and 2.5 μm (h).

**Figure 4.**
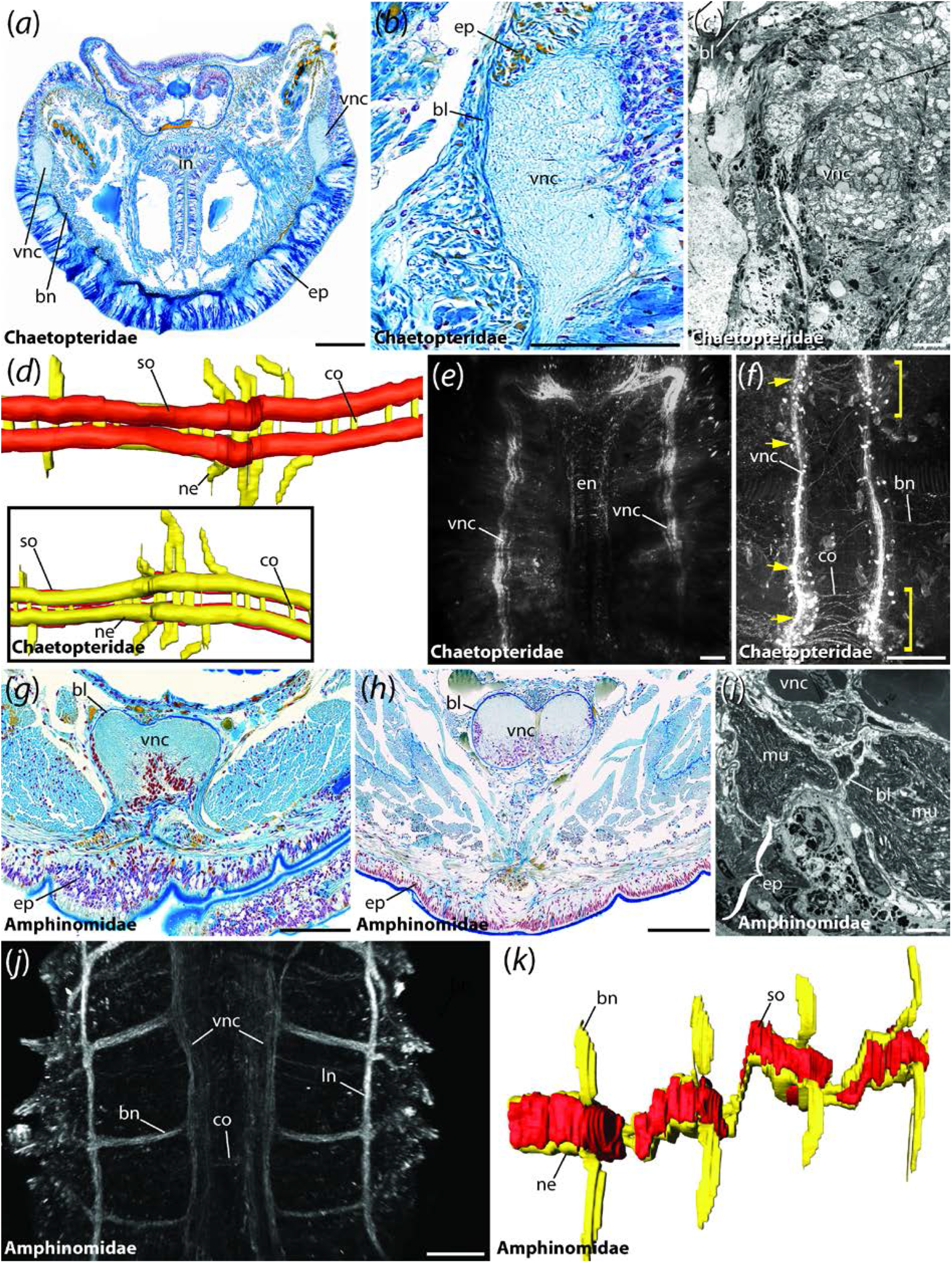
The VNC of Chaetopteridae and Amphinomidae. Cross-sections (a-c, g-i), ventral (d-f, j, k) and dorsal view (d, inset) of the ventral cord in *Spiochaetopterus costarum* (a-c), *Chaetopterus variopedatus* (d), *Phyllochaetopterus* sp. (e, f), *Eurythoe complanata* (g, h, j, k) and *Paramphimone* sp. (i). Anterior is up in (e, f, j), right in (d) and left in (k). Azan staining (a, b, g, h), TEM (c, i), immuno-staining against 5-HT (e, f) and tubulin (j), and 3D-reconstruction of serial histological sections (d, k). (a, b) The VNC (vnc) in Chaetopteridae is situated within the epidermis (ep). The basal lamina (bl) demarcates the epidermal from the subepidermal tissue. Giant fibers are absent. (c) Ultrastructural data of the same area reveal a position of the ventral nerve cord (vnc) within the epidermis (ep). A basal lamina (bl) is present underlying the epidermis (ep). (d) 3-D reconstruction of the posterior (vnc) reveals somata located along the entire ventral nerve cord and serial arrangement of the main commissures (co). (e) 5-HT labeling of the anterior body shows two parallel neurite bundles forming the ventral nerve cord (vnc). The esophageal nerves (en) are visible as well. (f) 5-HT labeling of the posterior body shows two parallel neurite bundles forming the ventral nerve cord (vnc) equipped with immunoreactive somata and serially and non-serially arranged commissures (co, also marked by yellow arrowheads) along the entire length of the cord. Note that the somata aggregations are marked by yellow brackets. (g) The anterior ventral nerve cord (vnc) in Amphinomidae is located within the epidermis (ep). A basal lamina (bl) demarcating the epidermal tissue is clearly visible. (h) Within the posterior parts of the body, the ventral nerve cord (vnc) is located in subepidermal position, outside the epidermis (ep). Nevertheless, a basal lamina (bl) interconnecting the (vnc) and the (ep) is present. (i) Ultrastructural data of the posterior ventral nerve cord (vnc) verify the presence of a basal lamina (bl) interconnecting epidermis (ep) and (vnc). (j) Anti-tubulin staining of the anterior ventral nerve cord (vnc) reveals the presence of branching nerves (bn), longitudinal nerves (ln) next to the (vnc) and commissures (co) along the (vnc). (k) 3-D reconstruction of serial histological sections supports the presence of hemiganglia containing the somata (so) and somata-free connectives along the (vnc) in Amphinomidae. bl, basal lamina; bn, branching nerve; co, commissure; ep, epidermis; in, intestine; ln, longitudinal nerve; mu, muscle; ne, neuropil; so, somata; vnc, ventral nerve cord. Scale bars = 100 μm (a, b, e-h, j), 5 μm (c), and 2.5 μm (j).

### Organization of the VNC in Palaeoannelida

Immunostainings against serotonin (5-HT) and FMRFamide as well as histological serial sections with subsequent 3D-reconstruction reveal a paired VNC within the first chaetiger in adult Oweniidae (figures 2a, d, f), whereas both neurite bundles form an unpaired mid-ventral cord containing a single neuropil in trunk chaetigers (figures 2b, d, f). Nevertheless, a bilaterally organized pair of neurite bundles showing certain immunoreactivity is detectable within this unpaired cord (figures 2d, f). An additional median neurite bundle is absent (figure 2b).

5-HT-LIR is present throughout the entire VNC (figures 2d, f). Notably, the somata are monomorphic (based on immunohistochemistry, histology and TEM), and scattered randomly along the ventral cord. The latter somata do not form distinct clusters - a result also supported by DAPI staining (figure 2f, inset) and histology (figure 2c). However, developmental studies in *Owenia fusiformis* reveal seriality of repeated 5-HT-LIR somata (figure 2d) [38] and investigations in adult *Galathowenia oculata* exhibit posterior somata clusters showing selective immunoreactivity [37]. Nevertheless, the latter results are not in contrast to the herein presented observations; they, illustrate the developmental complexity on the one hand, but also the necessity of a multi-methodological approach on the other hand, as well as the importance of future analyses of larval and juvenile stages in Oweniidae. Taken together, ganglia as defined by Richter et al. [62] were not observed.

In Magelonidae, immunohistochemistry reveals a paired VNC consisting of prominent neurite bundles and paired neuropils in thoracic chaetigerous segments (figures 2i, j) and a fused ventral cord in the trunk of adults (figure 2k). An unpaired bundle of median neurites described previously [65] could not be observed. Depending on the species both ventral neurite bundles may fuse between the ninth and tenth chaetiger (figure 2k). Nevertheless, neurites showing certain immunoreactivity still form paired neurite bundles. 5-HT-LIR (FMRFamide-LIR) is detectable throughout the ventral cord and – together with histology - revealed a non-serial arrangement of somata in the trunk. Somata-free connectives are absent (figure 2j, k). Irregularly arranged commissures not following a strict serial pattern are present (figures 2j), any seriality is missing.

Additionally, giant nerve fibers are present in all investigated magelonid species (figure 2h). Two parallel giant fibers originate in the brain, encircle the mouth and form a mid-ventral fiber, which runs along the trunk ventral in the main cord (figure 2k). To which extent the giant fibers are formed by one single or multiple axons cannot be stated based on the current dataset. However, giant fibers are lacking in most oweniids (figures 2a, b), but are found in *Myriowenia* sp. (figure 2b (inset)) where two parallel giant fibers fuse posterior of the mouth (not shown) and run along the entire trunk dorsal in the main nerve cord.

### Parallel neurite bundles in Chaetopteriformia

In Chaetopteriformia two parallel ventral neurite bundles are present in adult specimens (figures 3c, I; 4d). Although these bundles were observed as being separated anteriorly and converge in the posterior trunk, at least in Chaetopteridae, they never form a single mid-ventral neuropil. Serial clusters of monomorphic somata with 5-HT- and FMRFamide-LIR are present, but somata-free connectives are absent at least in Apistobranchidae and Chaetopteridae - a fact supported by histology (figures 3c, d; 4d-f). Serially repeated commissures interconnect both cords in Apistobranchidae (figures 3c, d), whereas mainly serial commissures as well as randomly arranged ones were seen in Chaetopteridae (figures 4d, f). In Psammodrilidae the commissures are serially arranged (figures 3e, g, i). In contrast to previous descriptions [41], but based on a comparable and even extended methodological approach, an unpaired midline neuropil is not detectable. Both, Apistobranchidae and Psammodrilidae, show a high degree of neuronal seriality, with only a few additional somata between the serial clusters of somata present in Psammodrilidae (figures 3e, g, f). Nevertheless, the nerve cord is intraepidermal in Chaetopteriformia.

### Occurrence of subepidermal VNCs in Amphinomidae and Sipuncula

In Amphinomidae, the trunk nervous system is represented by paired mid-ventral neurite bundles and additional bilateral longitudinal neurite bundles [35]. In the following, solely the median-most nerve fibers are regarded as part of the ventral cord and the lateral longitudinal ones, which are often described to belong to the peripheral nervous system, are not considered here. Thus, tubulin-immunoreactivity showed two ventral neurite bundles throughout the trunk (figure 4j). In anterior histological sections these paired neurite bundles form a (partly) fused mid-ventral cord in intraepidermal position and separate neurite bundles were only visible using immunohistochemistry (figure 4g). Posterior histological sections reveal two parallel neurite bundles forming the subepidermal cord (figure 4h), which, however, is still connected to the epidermis by a continuous ECM. A median neurite bundle and giant fibers are absent (figures 4g, h). The somata are clustered in certain parts of the neuropil and form segmental (hemi-) ganglia (figure 4k) in accordance with earlier descriptions [35], somata between the clusters and giant fibers are absent. Contrary, adult Sipuncula possess an unpaired medullary ventral cord with subepidermal somata and neuropil [69, 70] (see also figure 7). Giant fibers are absent.

A limited number of studies dealing with nervous system development in Annelida (including Sipuncula) shows that the subepidermal annelid VNC develops from a larval intraepidermal ventral cord [22, 69, 71]. Our analyses (figures 6, 7) also support a partial transition of the VNC from an intraepidermal towards a subepidermal position in the common stem lineage of Amphinomidae and Sipuncula, and a complete transition of the VNC towards a subepidermal position along the branch leading to Sipuncula.

A shift towards a subepidermal localization could also be observed in several Pleistoannelida (figure 7; see also figure 5 for Sabellariidae). On the other hand, numerous pleistoannelid groups, e.g., Polygordiidae [66], Tomopteridae (figure 5) or Siboglinidae [67, 68] and several other taxa bear a medullary-like VNC in intraepidermal position [21] (figure 7). However, many annelid groups are poorly investigated and developmental studies are necessary to back-up the findings observed in the adult ventral cord.

**Figure 5.**
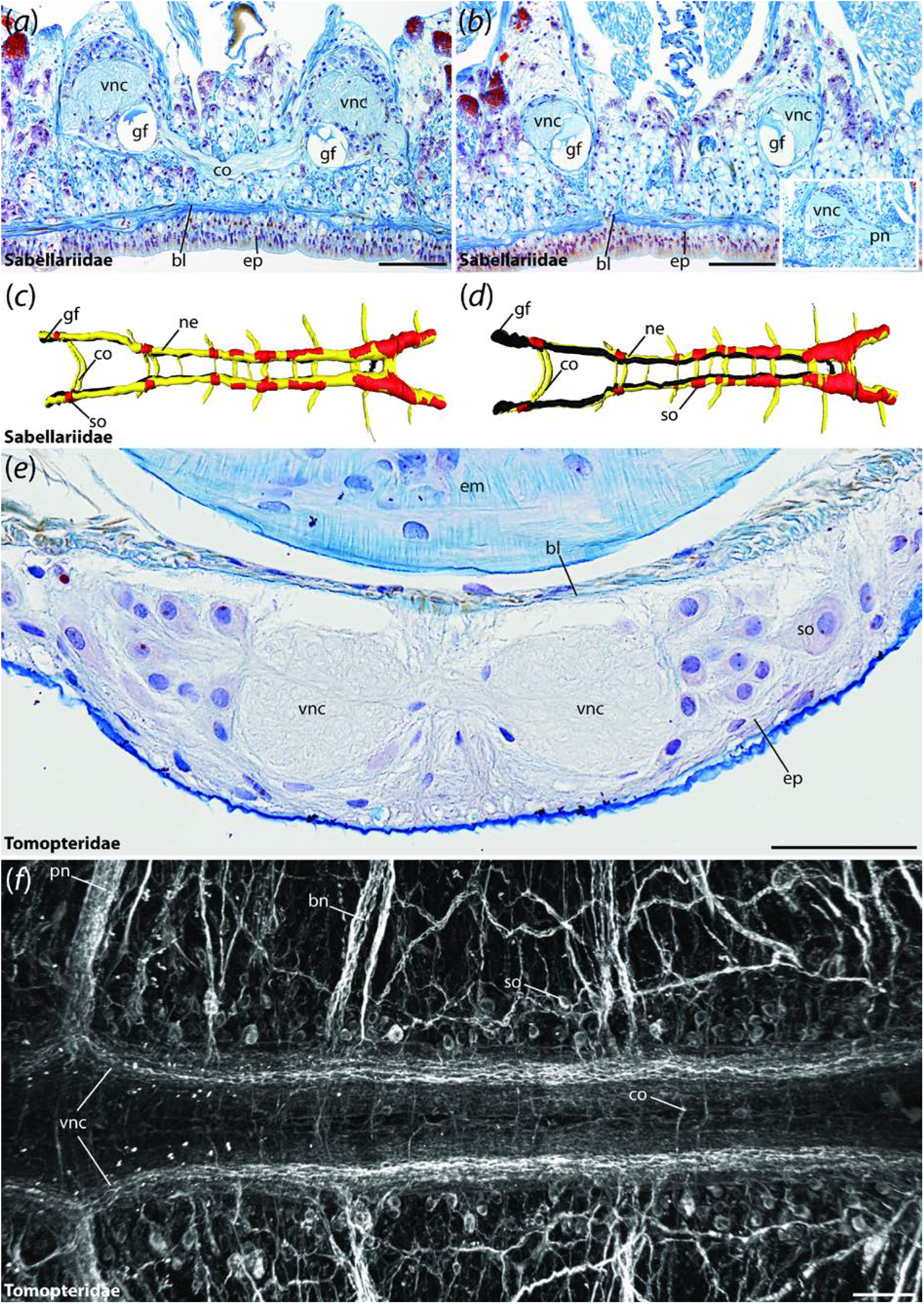
The VNC of Sabellariidae and Tomopteridae. Cross-sections (a, b, e) and dorsal (c) and ventral view (d, f) of the ventral cord in *Sabellaria alveolata* (a-d) and *Tomopteris helgolandica* (e, f). Anterior is right in (c, d, f). Azan staining (a, b, e), 3D-reconstruction (c, d) and immunostaining against α-tubulin (f). (a, b) The ventral nerve cord (vn) in *Sabellaria* is situated in subepidermal position. The ventrally located giant fibers (gf) are visible throughout the entire length of the animal and numerous serially arranged commissures (co) are present. The inset shows an outgoing parapodial nerve (pn) branching of from the ventral nerve cord. (c, d) 3D-reconstructions reveal the presence of serial somata (so), bearing ganglia and commissures (co). (e) The ventral nerve cord (vnc) in *Tomopteris* is situated within the epidermis (ep) and consists of two distinct neurite bundles. (f) Immunohistochemistry reveals the presence a medullary-like arrangement of somata (so), serially arranged commissures (co) and branching nerves (bn). Somata-free connectives are absent. bl, basal lamina; bn, branching nerve; co, commissure; em, esophageal musculature; ep, epidermis; gf, giant fiber; ne, neuropil; pn, parapodial nerve; so, somata; vnc, ventral nerve cord. Scale bars = 200 μm (a, b) and 100 μm (e, f).

**Figure 6.**
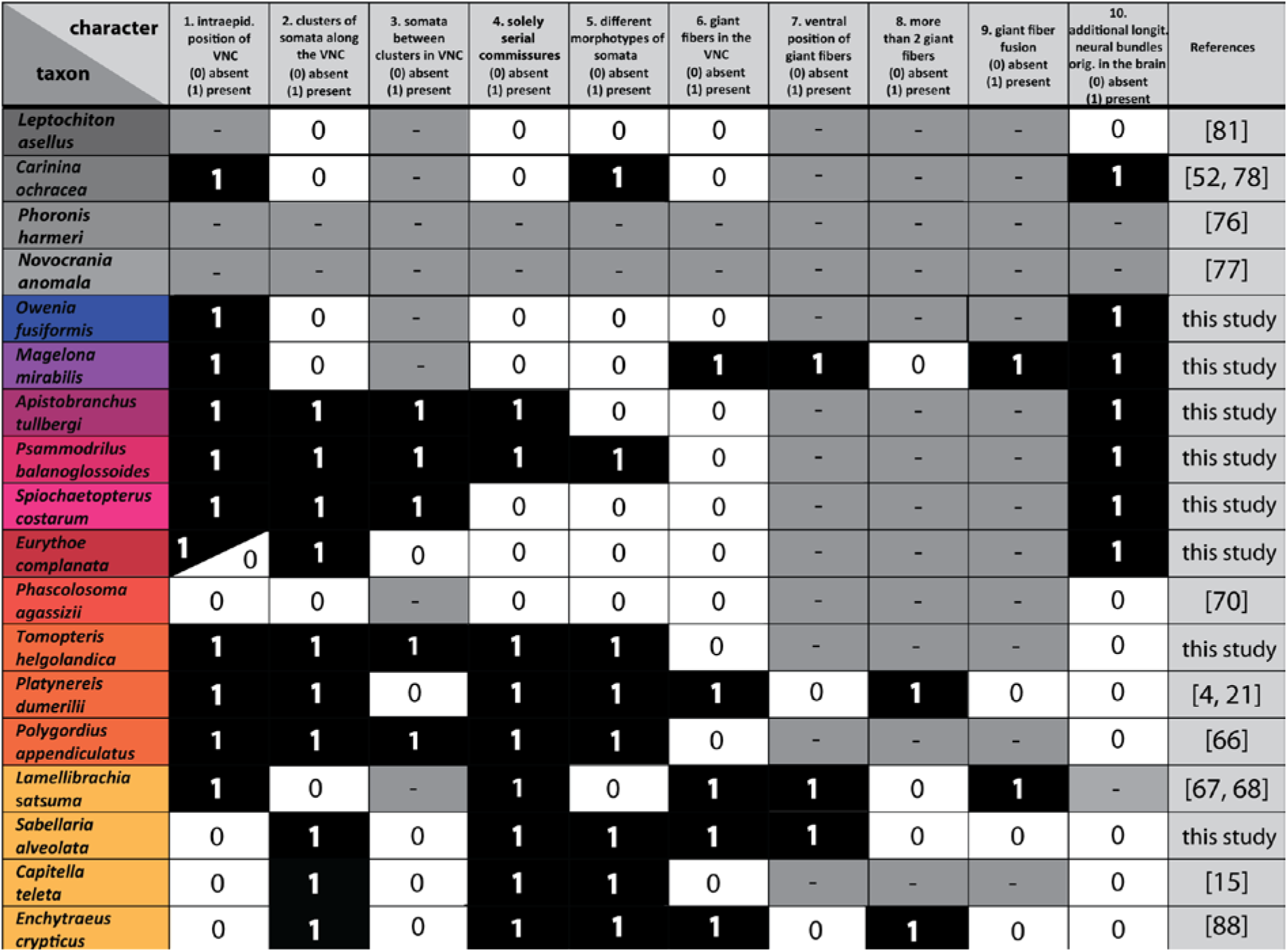
The morphological data matrix underlying the reconstruction of the ancestral state. Characters are coded as following: absent (0, white), present (1, black) and inapplicable (-, grey). References are included where required. When no reference is given, the data were raised during this study.

**Figure 7.**
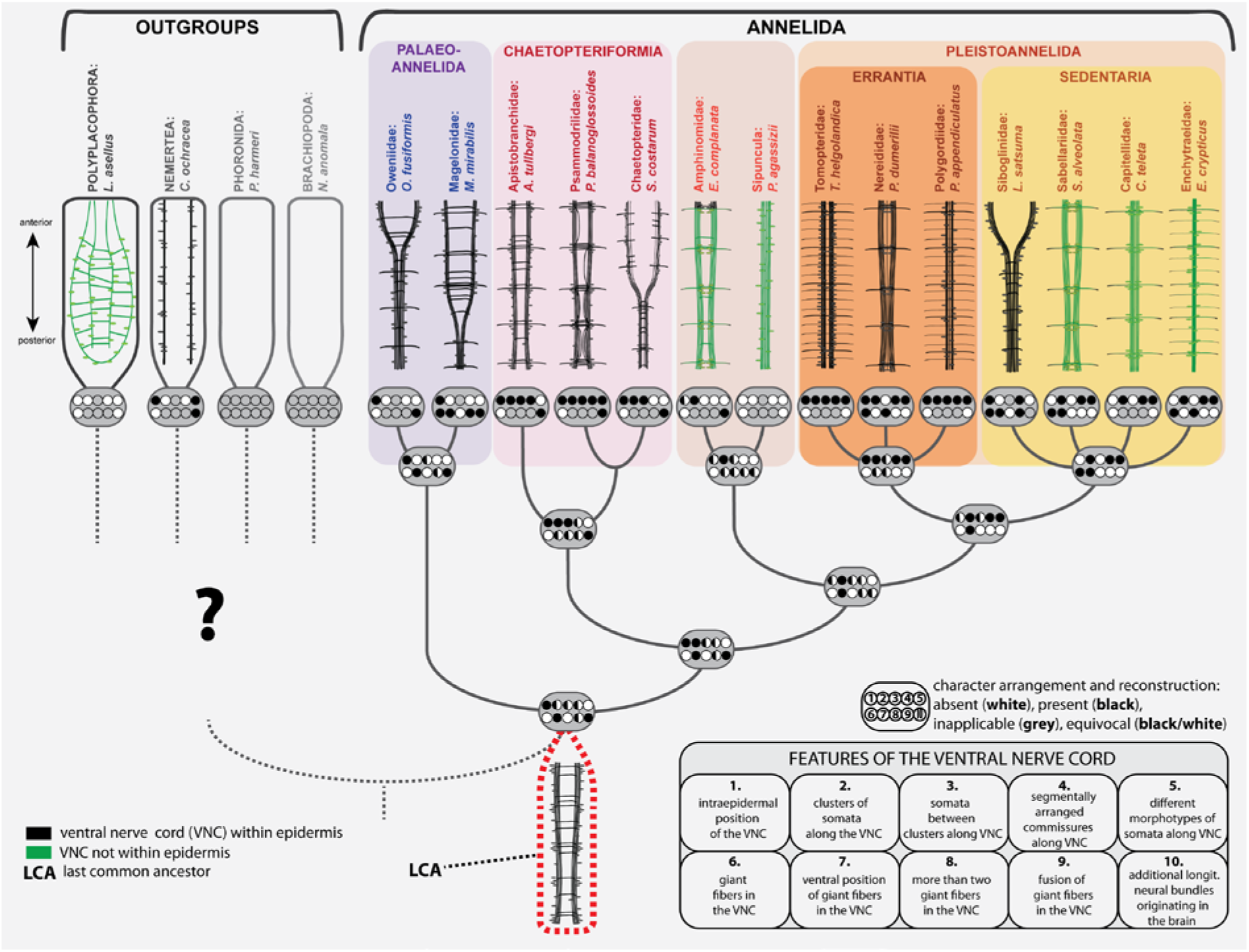
Schematic overview of the neural anatomy concerning the VNC in selected non-annelid taxa, the basally branching annelids and representatives of the Pleistoannelida. The shown topology is simplified and based on the MARE1.5 dataset. The character states given at the nodes of the topology are based on ancestral state reconstruction using maximum parsimony and the Mk1 model implemented in MESQUITE for the 10 characters specified for the VNC (electronic supplementary material S4-S5). Note that the question mark for the position of outgroups is caused by uncertain phylogenetic placements of the latter. Characters mapped on the topology are coded as following: absent (white), present (black) and inapplicable (grey). In cases when different character states were calculated for one character, all states are shown. Naming of each character is given in the figure.

### Nervous system evolution within Annelida

Our analyses using the MARE1.5 topology reconstruct an intraepidermal position of the VNC equipped with monomorphic somata to be the ancestral condition in Annelida (figure 7). Whether medullary arranged somata and non-segmental commissures were features of the last common annelid ancestor cannot be reconstructed unambiguously based on the current data and is highly dependent on the potential annelid sister group (figures 7; S6-S9). Nevertheless, the last common ancestor of Pleistoannelida most likely also had an intraepidermal nerve cord, but with somata clusters, intermediate somata and different morphotypes of somata. Moreover, giant fibers did not occur in the last common ancestor according to the reconstruction based on maximum parsimony (nevertheless equivocal with with ML ancestral state reconstruction). However, the existence of giant fibers in at least one member of the early branching Oweniidae and in all investigated Magelonidae, but lack of such neuronal structures in the remaining early branching lineages, makes it currently difficult to reconstruct the evolution of this character unambiguously (figure 7). Furthermore, their characterization mainly by diameter [92] and their variable location relative to the VNC when comparing Paleo-to Pleistoannelida add to this problem. Nevertheless, presence of such structures in members of Paleoannelida provides some support in the ML ancestral state reconstruction (additional file figure S5) to the scenario that giant fibers could belong to the annelid ground pattern. Due to the occurrence of giant fibers in only one oweniid species and the unresolved placement of the respective taxon within the oweniid tree, a final statement regarding the evolution of the giant fibers is hardly possible based on the data available.

Based on our analyses, a transition of the VNC into a ladder-like, cluster-bearing VNC with a subepidermal position seem to have evolved independently in a few lineages after the split of Pleistoannelida into Errantia and Sedentaria. A ladder-like, cluster-bearing, subepidermal VNC therefore has to be considered the derived condition within Annelida. The features of the VNC we predict for the annelid ground pattern are therefore differing from the commonly accepted, strict subepidermal ladder-like configuration (meaning somata only within the paired ganglia, paired somata-free connectives and serial commissures). This strict configuration has often been used to picture the ancestral annelid ventral nervous system and was largely based on the erroneous interpretation of the clitellate VNC as being the best fitting model for representing the annelid organization [1, 2, 21]. Nevertheless, a ladder-like but intraepidermal cord can be observed within Chaetopteriformia (in Psammodrilidae and partly in Apistobranchidae) as well as in several Pleistoannelida, and therefore it has to be assumed that the transition into this ladder-like condition occurred several times during annelid evolution. These presumably multiple transitions into a ladder-like appearance seem to show an evolution independent from the observed transition from an intra-towards a subepidermal position of the cord.

Notably, the position of the VNC is correlated with the arrangement of the body wall musculature. Generally, the longitudinal musculature is arranged in 2-3 pairs of bundles, in the ventral midline often separated by the VNC [85–87]. A body wall musculature consisting of inner longitudinal and dense outer circular muscle, as described for, e.g., burrowing Arenicolidae or members of Clitellata, was always thought to be part of the annelid ground pattern but is only present in a limited number of taxa. In addition, these are usually found in highly derived positions in the phylogenetic tree. Yet, such a body plan of ground-dwelling annelids is now found to be highly derived [22] and the subepidermal position of their ventral nerve cord may either be a protective adaptation to the mechanic stress during burrowing or related to the necessity of a well-developed system of circular muscle fibers. Notably, in species with an intraepidermal VNC, the ventral longitudinal muscle bundles are usually separated from each other by the VNC and true circular musculature forming closed rings of fibers is absent [86, 87].

A significant change in lifestyle, e.g., towards a more actively burrowing behavior in sediments like soil as depicted by numerous sedentary worms, or the exploitation of new food sources as seen, e.g., in predatory leeches, might have caused adaptive changes of the body wall musculature to fulfill specific movements required for this lifestyle. This might have triggered the necessity of a better protection of the nervous system, which caused a positional shift of the VNC. The latter or a similar scenario might help to explain the positional changes observable in the annelid VNC and is supported by comparable hypotheses dealing with the evolutionary transition of the brain in sediment-dwelling taxa towards posterior caused by the burrowing lifestyle [88]. Nevertheless, detailed investigations are necessary to point out the actual driving force for this major morphological change during annelid and spiralian evolution.

### Considerations about nervous system evolution within Spiralia

The evolution of the centralized nervous system within Bilateria is still a highly discussed field, and even similarities in terms of development and patterning of the ventral nerve cord in Spiralia are questioned in recent analyses – also due to lack of morphological data and comparable characters of the ventral nerve cord in various taxa [90, 91].

Although our results support an intra- or basiepidermal VNC and show the equivocal probability of somata occurring along the entire nerve cord as being the plesiomorphic annelid condition, the origin of the latter character state within Spiralia can hardly be reconstructed since the sister group of annelids is still not resolved [72, 73] (see also S6-S9 for ancestral state reconstructions including the potential sister group). Whereas either Nemertea or Mollusca were generally considered the annelid sister taxon [74, 75], recent analyses reveal Brachiopoda and Phoronida as additional candidates [31, 73]. In the latter, adults lack a VNC [6, 76, 77], but the nervous tissue within the lophophore is well developed and has an intraepidermal position as well. When using Brachiopoda/Phoronida as annelid sister group in the ancestral state reconstruction (MP analysis), the ground pattern of the annelid VNC includes an intraepidermal position of the VNC, monomorphic somata, a ventral position of giant fibers and additional longitudinal neural bundles. Nevertheless, presence of giant fibers in general and more than two giant fibers is not supported by this analysis. For the remaining characters an equivocal probability is given based on the MP reconstruction (figure S8/9). Nemertea also miss a VNC, but bear lateral medullary neurite bundles without strict somata clusters [78]. Both lateral neurite bundles are intraepidermal in the supposedly early branching Carininidae [52]. When using Nemertea as potential sister group for Annelida, the MP ancestral state reconstruction favors an intraepidermal position of the annelid VNC, a ventral position of giant fibers and the presence of additional longitudinal neurites as well. Somata clusters, segmentally arranged commissures, different somata morphotypes and giant fibers in general might be absent in the annelid ground plan based on this analysis (figure S7). Mollusca show a considerable variation in nervous system organizations, but the proposed mollusk ground pattern comprises a subepidermal nervous system out of four medullary cords, which are laterally and ventro-laterally positioned [79–81]. An inclusion of Mollusca into the analysis supports the lack of somata clusters, serial commissures and different somata morphotypes, but an intraepidermal position is questioned due to the subepidermal conditions of the VNC in molluscs. (figure S6) Yet, a subepidermal cord with distinctive somata clusters containing serially arranged commissures and with somata-free connectives in between is not depicted as the ancestral annelid configuration in none of the previously mentioned analyses (figure 7). Notably - as discussed above - such a configuration is also lacking in the supposed annelid sister groups [76, 77, 78, 81].

In numerous annelids the neurite bundles of the VNC are located ventro-laterally in early developmental stages (and in several lineages of interstitial or meiofaunal annelids), but mid-ventrally in adults [21, 68, 82–84]. Regardless of the annelid sister group a positional shift of the lateral neurite bundles towards the ventral midline has to be assumed to have evolved in the stem lineage of Annelida. According to our tree a similar transition must be hypothesized for the position of the VNC in Annelida. Here, the developmental shift of the annelid VNC from an intra-into a subepidermal position provides an explanation for the repeated evolutionary transition of VNC from the epidermis into deeper layers.

### Conclusions

Taken together, our study illustrates the complexity when it comes to evolutionary changes of organ system morphologies, but also shows the importance of phylogenetic analyses to test alternative hypotheses, e.g. regarding the direction of evolution. Based on the presented data an intraepidermal ventral nerve cord not only exhibits a larval and juvenile character as described previously for many annelid groups, but also reflects the plesiomorphic annelid condition. Accordingly, this condition is not (only) the result of paedomorphosis, as it was supposed for various annelid groups. Furthermore, profound scenarios concerning the evolutionary direction of changes in a respective organ system are solely possible under consideration of a comprehensive methodological approach, but also strongly limited by insufficient comparable datasets. In case of the ventral nerve cord in Annelida, further anatomical investigations are necessary to provide a better taxon sampling and data acquisition especially within the pleistoannelid groups. Only based on additional analyses open questions such as the evolution of giant fibers or the VNC transition from intra-towards subepidermal within Pleistoannelida can be resolved adequately. Thus the current study thereby provides important starting points for future investigations.

## Declarations

### Ethics approval and consent to participate

Ethics approval and consent to participate were not required for this work.

### Consent for publication

Not applicable.

### Availability of data and material

All data generated or analysed during this study are included in this published article and its supplementary information files. The assembled molecular datasets underlying the phylogenetic analyses are available in the Dryad digital repository; http://dx.doi.org/xxxxx [89].

### Authors’ contributions

CH and CB conceived the study, analyzed the data and drafted the manuscript. AW, THS and CB performed molecular analyses. CH, PB, TB, SHD, IK, GP and KW performed morphological investigations or data analyzes and contributed specimens or raw data. All authors discussed, read and approved the final version of the manuscript.

## Acknowledgments

We thank the Sars International Centre for Marine Molecular Biology (Bergen/ Norway), in particular H. Hausen for discussions and specimens of *Apistobranchus tullbergi* for TEM and the team of S11 for continuous support. We thank Pat Hutchings (Sydney/ Australia) for specimens of *Myriowenia*, Anett Karl (Leipzig, Germany) for TEM images of *Owenia fusiformis*, Nadya Rimskaya-Korsakova for helpful comments on the siboglinid nervous system and Helge Norf (Leipzig/ Germany) for help with the collection of *Hypania invalida*. Furthermore, we would like to thank the Arctic Station Greenland (Qeqertarsuaq, Disko, Greenland) and the biological station Roscoff (Roscoff, France), in particular Stephan Hourdez (Roscoff, France), for help and support during various collection trips.

## Additional files

**Additional file 1: Table S1.** Sampling sites and fixation/preservation details.

**Additional file 2: Table S2.** List of taxa used in the phylogenomic study and accession number. Species and accession numbers in bold were either newly sequenced or resequenced for deeper coverage in the present study.

**Additional file 3: Figure S3.** Best maximum likelihood (ML) tree of the RAxML analysis using the MARE2 data set of 40 taxa, including 404 gene partitions comprising 128,186 amino acid positions. Only bootstrap values above 50 are shown.

**Additional file 4: Figure S4.** Ancestral state reconstructions for the separate characters of the ventral nerve cord using a parsimony model with characters treated as unordered in MESQUITE v. 3.10. The character state is color coded and shown on the respective branch.

**Additional file 5: Figure S5.** Ancestral state reconstructions for the separate characters of the ventral nerve cord using the maximum likelihood Mk1 model with branch lengths scored as equal in MESQUITE v. 3.10. The character state is color coded and shown on the respective branch.

**Additional file 6: Figure S6.** Ancestral state reconstructions for the separate characters of the ventral nerve cord using a parsimony model with characters treated as unordered and Mollusca as outgroup in MESQUITE v. 3.10. The character state is color coded and shown on the respective branch.

**Additional file 7: Figure S7.** Ancestral state reconstructions for the separate characters of the ventral nerve cord using a parsimony model with characters treated as unordered and Nemertea as outgroup in MESQUITE v. 3.10. The character state is color coded and shown on the respective branch.

**Additional file 8: Figure S8.** Ancestral state reconstructions for the separate characters of the ventral nerve cord using a parsimony model with characters treated as unordered and Phoronida as outgroup in MESQUITE v. 3.10. The character state is color coded and shown on the respective branch.

**Additional file 9: Figure S9.** Ancestral state reconstructions for the separate characters of the ventral nerve cord using a parsimony model with characters treated as unordered and Brachiopoda as outgroup in MESQUITE v. 3.10. The character state is color coded and shown on the respective branch.

